# CD2 expression is co-regulated with stemness- and exhaustion-associated factors in human T cells

**DOI:** 10.1101/2025.01.10.632071

**Authors:** Philippos Demetriou, Maria Iakovou, Gregoria Gregoriou, Dimitris Vrachnos, Jianxiang Chi, Vasilia Tamamouna, Stavros Constantinou, Vakis Papanastasiou, Athos Antoniades, Paul Costeas

## Abstract

The CD2-CD58 pathway has been highlighted as a major player in anti-tumour T cell immunity. Our study reveals that CD2 costimulation strength significantly correlates with T cell activation, the average number of cell divisions, fold expansion, and IFN-γ production. Our findings suggest that the correlation of CD2 strength with the level of CD25 expression is a potential regulatory mechanism by which CD2 strength enhances above proliferation parameters. We find that human brain cancer tumour-infiltrating CD8+ and CD4+ T cells exhibit reduced levels of CD2, suggestive of a compromised CD2 strength upon CD2 engagement. Through a genome-wide CRISPR-Cas9 knockout screen, we identified two epigenetic regulators, SUZ12 and BAP1, as positive modulators of CD2 expression. We demonstrate that BAP1 is crucial for the upregulation and sustained high expression of CD2 following T cell activation. We reveal that CD2 is co-regulated with other co-stimulatory/inhibitory receptors, and factors associated with T cell stemness and exhaustion, in a dose-dependent manner. Importantly, we rescue the loss of CD2 due to BAP1 knockout by pharmacological inhibition of histone deacetylases making this a harnessable regulatory pathway. The insight from our study enhance our understanding of CD2-mediated T cell regulation and identify essential regulators of this pathway.

## INTRODUCTION

The interplay between costimulatory and coinhibitory receptors on T cells is crucial for determining downstream T cell activation and responses^1,2^. Ιmmunotherapy strategies aimed at enhancing T cell immunity against cancer have focused on either blocking coinhibitory receptor-ligand interactions or activating co-stimulatory receptor-ligand pathways. Despite these advances, our understanding of the functions and regulatory mechanisms of these co-signaling receptors remains incomplete. One such notable pathway is the one initiated by the interaction of CD2 receptor on T cells and CD58 ligand on target cells, which has garnered significant attention recently^2,3^. Patricia Ho *et al*^2^. and Beiping M. *et al*.^4,5^ identified CMTM6 as a positive regulator of CD58 in tumour cells. They found that competition for this regulator between CD58 and the coinhibitory ligand PD-L1, which binds to PD-1, plays a critical role in shaping downstream T cell anti-tumor responses.

We and others have demonstrated that peripheral blood CD4+ and CD8+ memory T cell subsets express higher levels of CD2 compared to their naïve counterparts^4,5^. We recently determined that the expression levels of CD2 on T cells influence the extent of CD2-CD58 interactions during T cell synapse formation^5^. These interactions, in turn, significantly amplify proximal T cell receptor (TCR) signaling. Furthermore, we observed that strong PD-1 signaling—characterized by elevated PD-1 expression akin to that found in tumor-infiltrating T cells (TI-T cells)—partially inhibits the CD2-mediated enhancement of proximal TCR signaling. Notably, in a significant percentage of patients with colorectal (CRC), endometrial (EndoC), and ovarian cancers (OC), we detected a tumour infiltrating (TI) CD8+ T cell population with significantly reduced CD2 expression, which we termed the CD2^low^ phenotype.

TI-T cells are frequently described as exhausted due to chronic antigen stimulation, characterized by a progressive loss of effector functions^6,7^. The previously recognized pool of exhausted TI-T cells has been further classified into progenitor exhausted (Tpex), transitory, and terminally exhausted (Tex) T cells^8,9^. This classification reflects their potential differentiation trajectory, with Tpex maintaining greater stemness properties than Tex cells, such as self-renewal, long-term maintenance and effector exhausted T cells capable of efficient tumour clearance. A Tpex signature has been associated with improved responses to immune checkpoint blockade and enhanced efficacy of adoptively transferred T cells against cancer^10–13^

Here, we employed ex vivo profiling of human T cells, in vitro functional assays, and genetic screening approaches to deepen our understanding of CD2 function and regulation. We determined that varying strengths of CD2 costimulation significantly affect downstream T cell proliferation and IFN-γ production. Additionally, we found that the CD2^low^ phenotype is also exhibited by glioblastoma and meningioma CD4+ TI-T cells alongside CD8+ T cells. In our quest to identify positive regulators of CD2 expression, we discovered that BRCA-1 associated protein 1 (BAP1) is essential for expressing and maintaining high levels of CD2 expression in T cells; moreover, we determined that BAP1 serves as a broader regulator of key co-signaling receptors involved in T cell function other than CD2 and factors regulating T cell stemness, and metabolism.

## RESULTS

### T cell proliferation and IFN-γ production correlate with CD2 costimulation strength

We previously demonstrated that interactions between CD2 and CD58 in the T cell synapse linearly amplify proximal T cell receptor (TCR) signaling. To investigate how the strength of CD2 costimulation affects downstream T cell responses, we monitored the proliferation of CFSE-labeled human peripheral blood (PB) naïve and memory T cells after polyclonal in vitro activation, with varying levels of CD2 costimulation and interleukin-2 (IL-2) cytokine. We observed that decreasing CD2 costimulation levels resulted in reduced percentages of proliferating cells, as shown in Fig. S1a-b. Focusing on the proliferating cells, we found that lower CD2 costimulation led to a significant decrease in both the average number of cell divisions and the fold expansion of these cells, as indicated by the proliferation index (PI) and replication index (RI), respectively; this trend was consistent across both PB CD4+ and CD8+ T cell compartments (Fig. 1a-b, S1c-d). Notably, PI and RI positively correlated with IL-2 receptor alpha (CD25) expression measured two days post-activation (Fig. 1c-d, S1e), suggesting that the strength of CD2 costimulation may regulate T cell proliferation following activation and IL-2 stimulation by modulating CD25 expression. A similar pattern was observed for IFN-γ production in an in vitro differentiation setting of naïve CD4+ T cells into Th1 cells. The effect mediated by CD2 was particularly pronounced for IFN-γ compared to IL-2 T cell production. A decrease in CD2 costimulation resulted in both a reduced percentage of IFN-γ-producing T cells and a lower level of IFN-γ produced by these cells, as illustrated by changes in high, intermediate, and low IFN-γ producers (Fig. 1e, S1f).

**Fig. 1.**
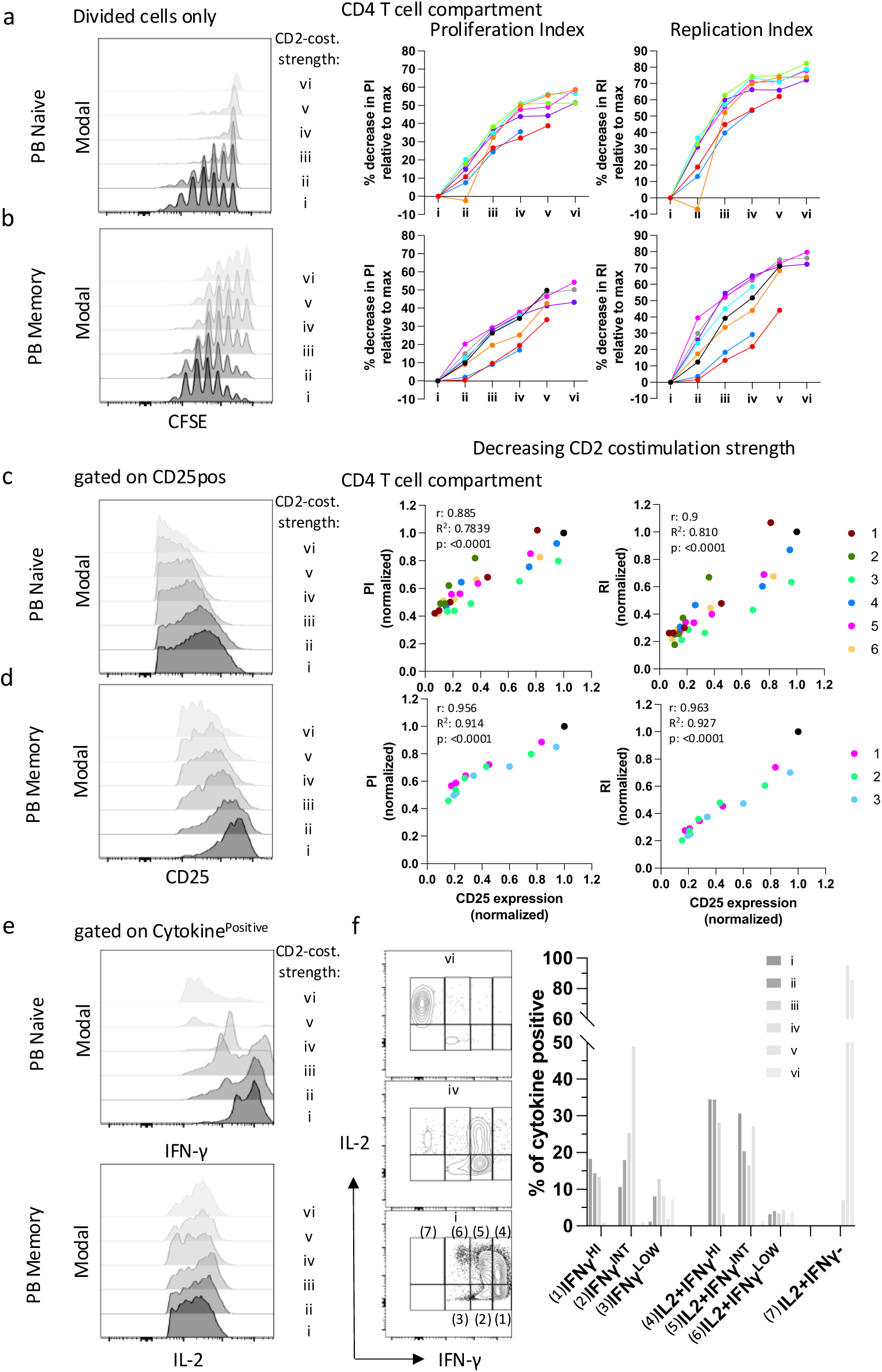
T cell proliferation and IFN-γ production correlate with CD2 costimulation strength. Representative flow cytometry histograms (left) from one healthy volunteer, illustrating cell divisions of PB CFSE-labelled naïve **(a)** and memory **(b)** CD4+ T cells, gated on divided cells and corresponding plots (right) of the percentage decrease in proliferation index and replication index at titrated levels of CD2 costimulation strength (i - vi), 5-days post *in-vitro* activation from 7 donors (each a different colour). From experiments as in (a), representative histograms (left) of CD25 expression in naïve **(c)** and memory **(d)** CD4+ T cells 2 days post *in-vitro* activation, and corresponding plots showing CD25 expression against PI or RI expression for 8 and 3 donors, respectively (all parameters normalized to value in highest CD2 costimulation strength condition (i) to allow pooling of all healthy donors samples in one plot). **(e)** Histograms of intracellular levels of IFN-γ (top) and IL-2 (bottom) expression of naïve CD4+ T cells differentiated to Th1 cells *in-vitro,* 7 days post-activation and **(f)** Quantification of percentages of 7 populations of cytokine producing subsets, as gated in representative IL-2 vs IFN-γ dot plots, in the presence of titrated levels of CD2 costimulation strength (i-vi); a representative donor shown from two independent healthy donor experiments.

### A CD2^low^ phenotype exhibited by brain tumour-infiltrating CD8+ and CD4+ T cells

We previously demonstrated that the expression levels of CD2 on T cells can limit CD2 costimulation, with PB memory T cells exhibiting a higher number of CD2 molecules compared to naïve T cells^5^. Building on these findings, we compared CD2 expression levels in T cells infiltrating human glioblastoma and meningioma (gli T and mng T cells) with their peripheral blood (PB) T cell counterparts. By categorizing PB, mng, and gli T cells using naïve and memory markers (CD62L and CD45RA) (Fig S2a), we observed reduced CD2 expression in both gli T and mng T cells within each memory-marker expressing (central memory, CM, effector memory, EM and EM re-expressing CD45RA, EMRA) CD8+ T cell compartment (Fig. 2a,c). Notably, this reduction aligned their CD2 expression more closely with that of corresponding naïve PB T cells across all patients; naïve T cells were almost completely absent from these tumours (Fig. S2a). This phenomenon is reminiscent of the CD2^low^ phenotype we previously described in colorectal (CRC), endometrial (EndoC) and ovarian cancer (OC) patients. In contrast to our earlier studies in CRC, EndoC, and OC cohorts, the CD2^low^ phenotype was also evident and strong in the tumor CD4+ T cell compartment for nearly all brain cancer patients (Fig. 2b-d). Our findings, expand our initial observation from other cancer types to brain tumours and reveal a unique difference in that the CD2^low^ phenotype is also exhibited by in gli and mng CD4+ T cells and not just gli and mng CD8+ T cells. This is important to consider since it has been shown that activation of both subsets is required for functional anti-tumour T cell immunity in brain tumours^14–16^

**Fig. 2.**
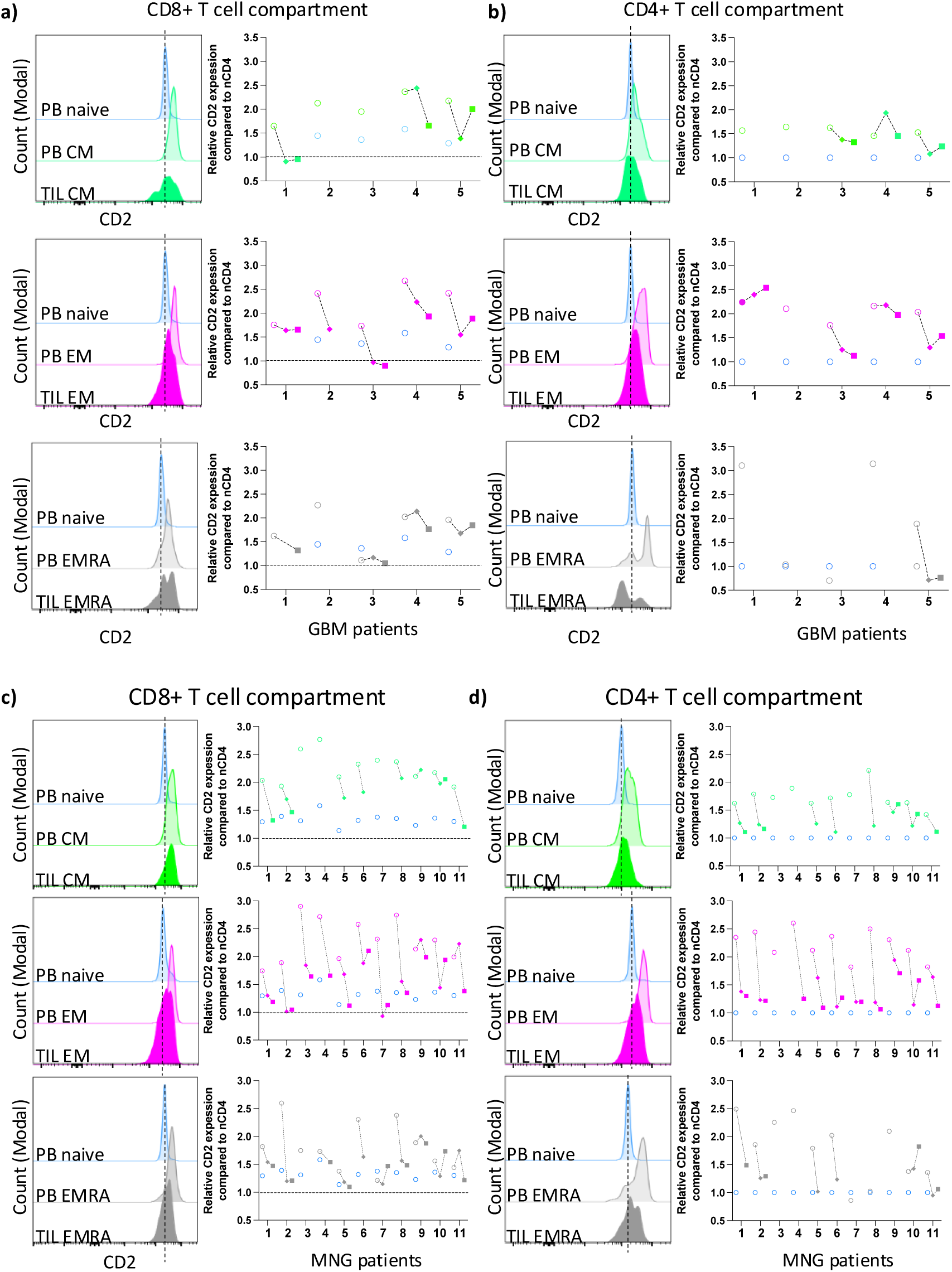
A CD2^low^ phenotype exhibited by brain tumour infiltrating CD8+ and CD4+ T cells. **(a-b)**, (left) Representative flow cytometry histogram of CD2 expression in naïve (blue), central memory (CM-green), effector memory (EM-mangenta) and effector-memory re-expressing CD45RA (EMRA-grey) CD8+ and CD4+ T cells, respectively, from peripheral blood (PB-light colour histograms) and tumour (solid colour histograms) from one glioblastoma patient. (**a-b),** (right). Relative CD2 expression in T cell subsets, normalized to CD2 expression in PB naïve CD4+ T cells, per donor per tissue as in (a-b, left) across 5 patients; open-light colour circles represent PB, solid rhombus core side of tumour and solid square peripheral side of tumour. **(c-d)**, Same as in (a-b) but for 11 meningioma patients.

### Genome-scale CRISPR-Cas9 screen identifies positive regulators of CD2

In light of the differential CD2 expression levels detected in the various T cell subsets, we sought to identify regulators of CD2. We performed a positive selection, using fluorescence-activated cell sorting (FACS), genome-wide CRISPR-Cas9 loss-of-function (LOF) screen in Jurkat T cells (Fig. 3a). Perturbations enriched in the positively selected T cell population with low CD2 surface protein (CD2lo) could be physiological positive regulators of CD2. Six targets were consistently enriched across different time points in the CD2lo T cell population compared to the control group, with CD2 itself being the most frequently identified target. (Fig. 3b-c, Supplementary Table 1). The nuclear deubiquitinating enzyme BRCA1-associated protein-1 (*BAP1*) that was shown to be essential for thymocyte development and proliferative responses of T cells, was the next most frequently enriched target after *CD2* hence our focus thereafter. Amongst the six enriched targets, there was another epigenetic regulator gene the SUZ12 polycomb repressive complex 2 (PRC2) subunit that forms part of the PRC2 repressive complex. Subsequent experiments involving independent BAP1 gene and SUZ12 knockouts (Kos) confirmed that BAP1 and SUZ12 regulate CD2 expression with the strongest effect seen upon BAP1 KO relative to SUZ12 loss (Fig. 4 d-f). Real-time qPCR indicated a reduction in *CD2* mRNA upon BAP1 or SUZ12 KO suggesting that they may exert their regulatory effect at the transcriptional level (Fig. 4g).

**Fig. 3.**
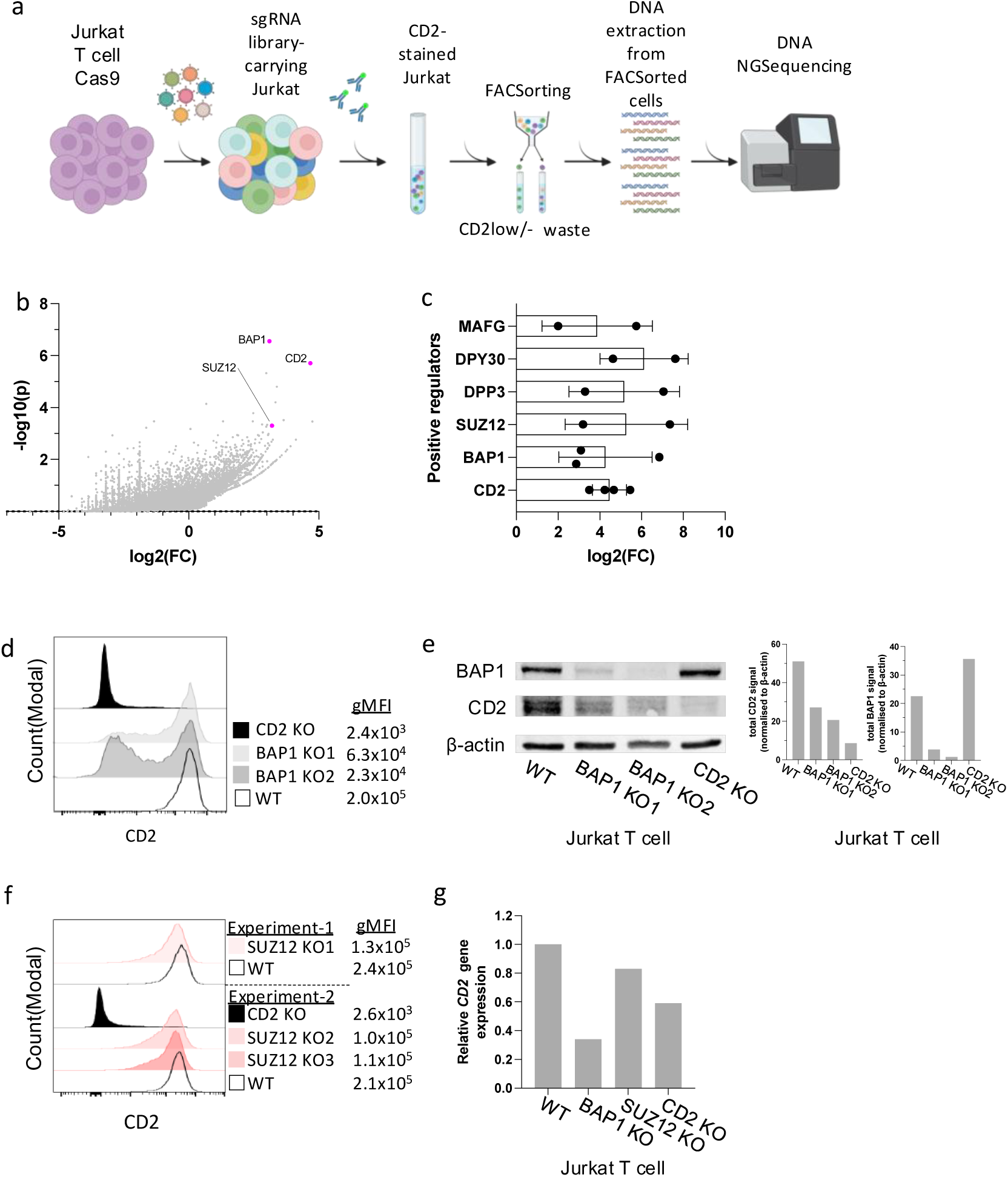
Genome-scale CRISPR-CAS9 screen identifies positive regulators of CD2. **(a**) Diagrams shows the experimental design used for identifying positive regulators of *CD2* using a whole genome-wide CRISPR-CAS9 KO screen in the human Jurkat T cell line. CD2lo expressing cells were FACsorted (step 3) at five different timepoints during the cell library screen culture. **(b)** A representative example plot from one of the timepoints, showing the distribution of target gene enrichment withing the CD2lo population compared to the total unsorted population based on log2 fold change of gene enrichment and p-value for enrichment generated by the use of the MAGeCK algorithm (REF). **(c)** Bar chart showing the log2 fold change enrichment of targeting sgRNAs in CD2lo against unsorted control population, that were reproduced in more than one of the 5 screen timepoints tested; each dot represents a separate timepoint. Data represent mean+/-SD. (**d)** Representative flow cytometry histograms of surface CD2 expression upon two different sgRNA targeting BAP1 for knock out (BAP1 KO1 and KO2) and one sgRNA targeting CD2 (CD2 KO) compared to WT control) and the corresponding geometric mean fluorescence intensity of each total population (gMFI) from two independent experiments. **(e)** Western blot and corresponding bar charts showing the total amount of BAP1 and CD2 protein normalized to beta-actin (b-actin) expression in experiments as in (d). **(f)** Histograms from two independent experiments are presented here, showing the surface CD2 expression upon three different sgRNA targeting SUZ12 for knock out (SUZ1 KO1, KO2 and KO3) compared to a CD2 KO and a WT control with corresponding geometric mean fluorescence intensity of each total population (gMFI). **(g)** Bar chart showing the relative *CD2* mRNA expression upon BAP1, SUZ12 and CD2 KO as in experiments (d-f) compared to WT control as quantified by real-time quantitative PCR.

**Fig. 4.**
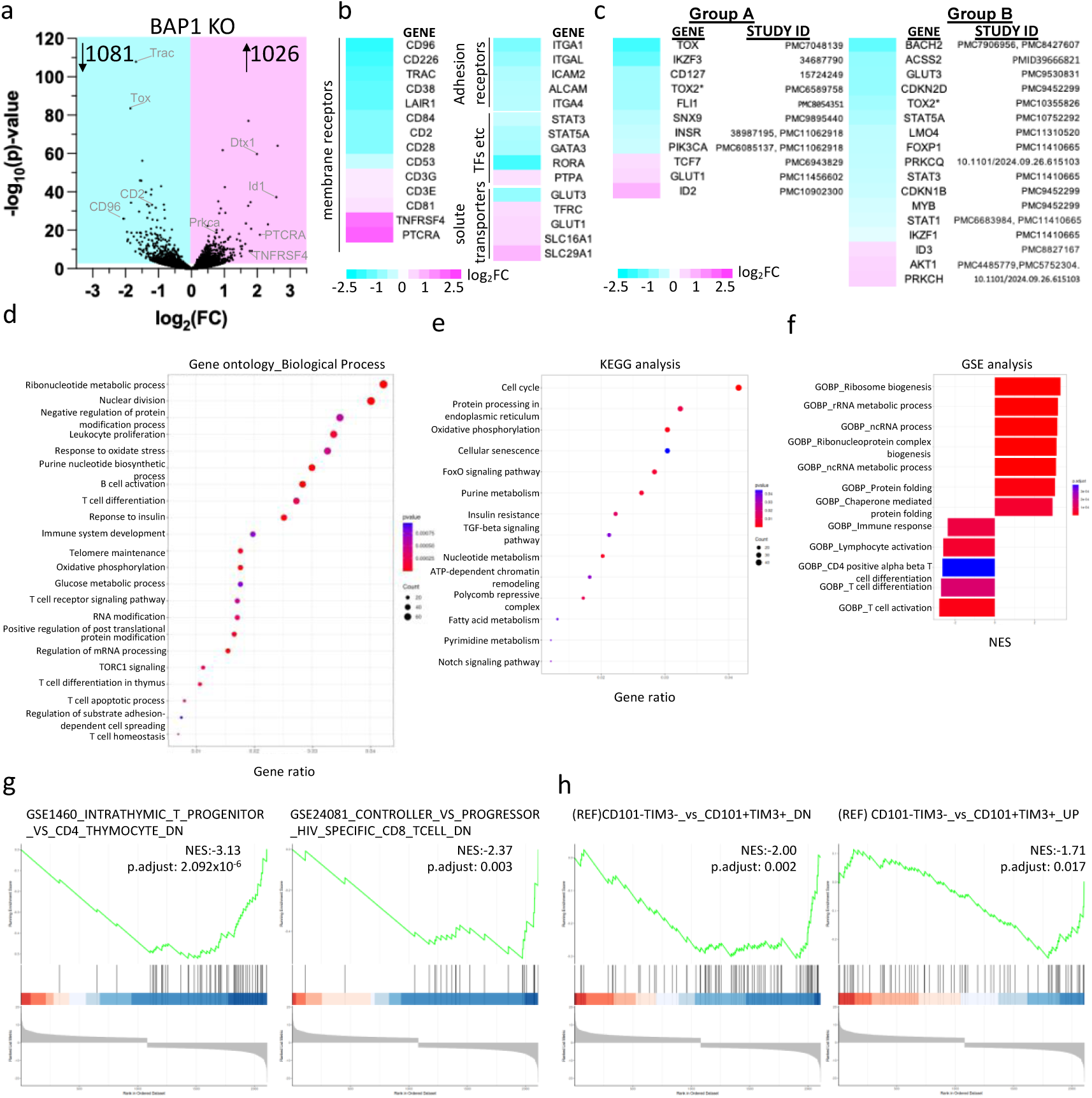
CD2 expression is coregulated with expression of markers associated with T cell development, stemness, exhaustion and metabolism. **(a)** Volcano plot showing p value and fold change in expression of genes detected in BAP1 KO Jurkats relative to WT Jurkats. **(b-c)** Heatmap of the log2 fold change (FC) expression of selected differentially expressed genes (DEGs) and their corresponding gene names identified in BAP1 KO Jurkats relative to WT control; these are categorized in T cell surface signaling molecules, adhesion molecules, transcription factors (TF) and solute transporters in (b) and in factors associated with stemness and exhaustion of T cells in (c). **(d-e)** Bubble plots showing a selection of enriched pathways determined by performing gene ontology analysis (biological process pathways) in (d) KEGG pathway analysis in (e) of DEGs in BAP1 KO T cells with a padjust<0.05. The gene count per pathway is represented by the size of the dot and the p value by the colour of the dot. (**f)** Bar chart showing a selection of enriched pathways and the corresponding normalised enrichment score (NES) determined by performing gene set enrichment analysis (GSEA) using the C5 genesets of the Molecular Signatures Database (MSigDB) and the same DEGs as in (d-e). **(g)** A selection of enriched gene signatures in BAP1 KO identified upon GSEA using the C7 immunologica signature gene sets in MSigDB. **(h)** GSEA in BAP1 KO T cells for a progenitor exhausted T cell gene signature generated from ^8^

### CD2 expression is co-regulated with expression of markers associated with T cell development, stemness, exhaustion and metabolism

To investigate whether BAP1 KO affects only CD2 expression expression, we conducted a bulk transcriptomic analysis comparing BAP1 KO Jurkat T cells to wild-type (WT) Jurkat T cells. BAP1 KO led to the identification of 2107 differentially expressed genes (DEGs) relative to the WT (Fig. 4a, log2FC > 0 and adjusted P-value < 0.05), with 1026 genes being upregulated and 1081 downregulated (Supplementary Table 2a). This altered transcriptional profile following BAP1 KO was characterized by the downregulation of key T cell surface receptors, beyond that of *CD2,* notably *CD28* and *CD226* with *TRAC* being amongst the top 10 DEGs, and the upregulation of others such as *CD3G*, *CD3E*, *TNFRSF4/OX40*, *PTCRA*, and *CD81* (Fig. 4b). Changes in total TRAC protein levels were validated using western blotting (Fig. S3a) and the loss of TCR complex at the cell surface and other surface receptors was confirmed using flow cytometry (Fig. S3b). Downregulated genes included molecules involved in adhesion and cell trafficking (*SELL*, *ITGA1*, *ITGA4*, *ITGAL*, *ICAM2*, *ALCAM*) (Fig 4b). These results indicate BAP1 regulation is not specific to CD2 but affects additional essential receptors involved in T cell development, T cell identity and migration. BAP1 KO resulted in alterations in genes regulating T cell differentiation such as RORA, GATA3, PTPA and solute transporters regulating metabolism such as GLUT1, GLUT3 (Fig. 4c). BAP1 KO resulted in DEG associated with T cell stemness, differentiation to memory, progenitor exhausted and exhausted T cells. Gene expression changes in group A genes (*TOX*, *TOX2*, *TCF7*, *ID2*, *SNX9* etc) (Fig. 4c) would favour T cell stemness based on separate studies investigating each individual gene independently. In contrast, gene expression changes in Group B genes (*ACSS2, ID3*, *MYB*, *PRKCQ*, *PRKCH*, *LMO4* etc) have been associated with reduced T cell stemness and would favour a T cell exhaustion state. TOX2 has been attributed contrasting roles with potentially context-dependent differences i.e. its expression imposes T cell exhaustion^17^ or promotes central memory differentiation and prevents exhaustion in a context dependent manner^18^. Interestingly, there was an increase in CD57+ T cells upon BAP1 KO CD57 expression and a 1.4-1.7-fold increase in CD57 average expression amongst CD57+ T cells compared to WT ones (Fig S4c). This is suggestive of increased terminal differentiation phenotype upon BAP1 loss as CD57 expression has been associated with terminally differentiated CD8+ T cells^19^ and T cell senescence^20^ but also with the responding T cells to anti-PD-1 checkpoint therapy in non-small cell lung cancer^21,22^.

Gene ontology analysis (Fig. 4d-e, Suppl, Fig. 3c, Supplementary Table 2b-c) confirmed BAP1’s role in regulating genes involved in T cell receptor signaling regulation, T cell synapse, adhesion processes, T cell differentiation, immune system development and metabolism such as glucose metabolism, and insulin response. It further revealed involvement of BAP1 in regulating genes involved oxidative phosphorylation, regulation of telomeres and post-translational protein modification pathways. KEGG analysis, revealed enrichment of pathways associated with cell cycle, Notch, TGF-beta and FoxO signaling pathways, nucleotide and fatty acid metabolism and finally oxidative phosphorylation and cellular senescence (Fig. 4e, Supplementary Table 2d).

Similar to the transcriptomic data findings, gene set enrichment analysis (GSEA) indicated a downregulation of processes related to immune responses, T cell activation and differentiation. In contrast, there was an upregulation of processes associated with ribosome biogenesis, non-coding RNA processing and metabolic processes (Fig. 4f, Supplementary Table 2e). GSEA also indicated similarities between BAP1 KO-induced gene changes and those observed in human intrathymic T progenitor cells compared to CD4 thymocytes, suggesting a role of BAP1 in thymocyte development. Additionally, parallers were drawn with gene expression differences in HIV-specific CD8 T cells of HIV controllers compared to progressors (Fig 4g, Supplementary Table 2f). Generating a Tpex gene signature by comparing the DEGs of Tpex (CD101-Tim3-T cells), to Tex (CD101+Tim3+ T cells) from a previous mouse study^8^ and in the context of mouse-human orthologs, we run a GSEA on this Tpex signature using our DEGs of BAP1KO T cells (Fig 4h). This revealed that BAP1KO T cells downregulate a number of genes also downregulated in Tpex. At the same time, BAP1KO T cells also downregulated genes that were found upregulated in Tpex. These findings, although contrasting between them, they agree with our manual interpretation of our transcriptomic findings (Fig. 4b) that show that upon BAP1 KO there are changes that can both favour and disfavor an enhanced T cell stemness state. In summary, under our experimental conditions CD2 was regulated together with a number of other costimulatory receptors and stemness-, exhaustion-associated factors.

### BAP1 has dose-dependent effect on costimulatory receptor expression and stemness-exhaustion transcription factors

Upon profiling T cells 7 days post sgRNA BAP1 transduction for BAP1 KO, we observed a significant increase in the percentage of the CD3-CD2+ subset (Single Positive, SP) compared to the control. Notably, it was not until 12 days post-transduction that a significant increase in the CD3-CD2-(Double Negative, DN) subset, indicating loss of CD2, was detected; this subset continued to rise over time, peaking around days 20 to 30 post-transduction (Fig. 5a and Fig. S4a). The expression profile of CD3 and CD2 achieved by the end of the experiment was maintained during further culturing of the T cells. Unexpectedly, BAP1 protein levels varied among these subsets, decreasing from SP to DN (Fig. 5b), with the highest expression found in the wild-type (WT) control, as expected. Similarly to the titrated levels of BAP1 observed in the DN and SP subsets, there was a corresponding titrated effect on the surface expression of co-stimulatory and adhesion receptors (Fig. 5c, Fig. S4b) shown to be downregulated at the protein level and upon bulkRNAseq analysis; this was observed using two different sgRNAs to KO BAP1 (Fig. 4b, Fig S4b). The DN subset exhibited the lowest levels of CD2, CD28, CD11a, and CD49a, with expression levels increasing in the SP subset and the highest being in the WT control; the same trend is observed in the percentage of positive cells for each marker. Notably, CD58 expression did not show significant differences between these subsets or compared to the WT control. A similar BAP1 dose-dependent effect on surface receptor expression was observed for transcription factor genes related to T cell stemness *(tox, tcf1, bach2, id2*) (Fig. 5d).

**Fig. 5.**
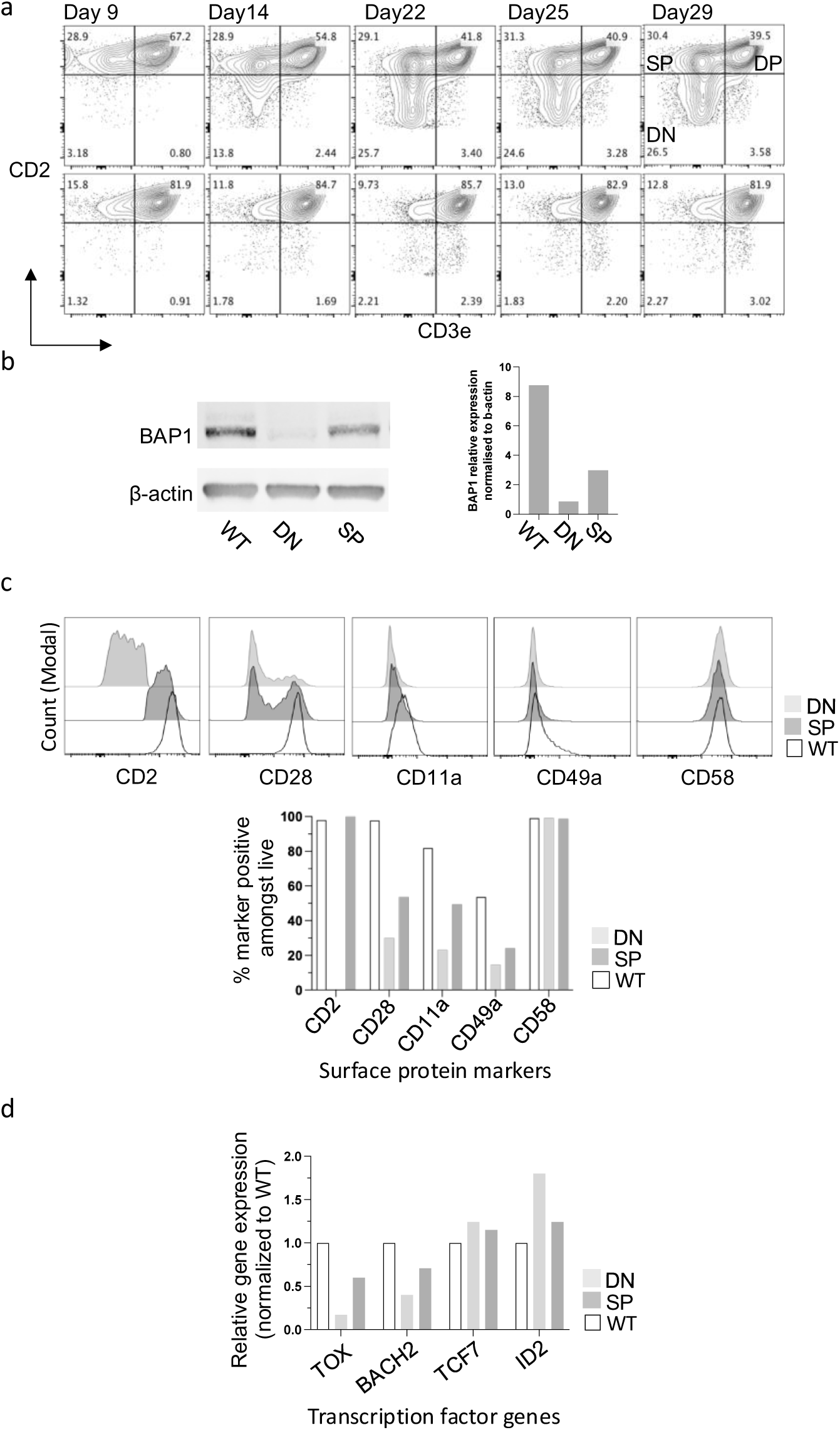
BAP1 has dose-dependent effect on costimulatory receptor expression and stemness-exhaustion transcription factors. **(a)** Flow cytometry contour plots showing CD2-CD3e expression profile during sgRNA BAP1 KO2 in Jurkat T cells at different timepoints of the culture and the gating to define different populations, double positive, DP: CD3+CD2+, single positive, SP: CD3-CD2+/high, double negative DN: CD3-CD2-/low. (**b)** Western blot and corresponding bar charts showing the total amount of BAP1protein normalized to beta-actin (b-actin) expression in experiments as in (a) at endpoint when culture showed no further deviation in ratio of DP, SP and DN subsets. **(c)** Flow cytometry histograms (top) of CD2, CD28, CD11a, CD49a, CD58 in DN, SP BAP1KO2 T cell subsets and WT T cells and the corresponding percentage positive T cells for each marker is shown in bar chart (bottom). **(d)** Bar chart showing the relative gene expression for stemness-exhaustion associated transcription factors in WT, single positive (SP) and double negative (DN) Jurkats. A reprentative profiling culture timepoint shown out of three.

### T cell activation reveals differential CD2 and TCR expression in WT versus BAP1 KO T cells but not for CD28 and CD226

CD2 expression is significantly upregulated following T cell activation. In DN BAP1KO T cells, the expression of the TCR/CD3 complex was nearly absent. To activate both wild-type (WT) and DN BAP1KO T cells, we employed phorbol myristate acetate (PMA) and/or ionomycin (ION) treatment at low and high concentrations and in combinations, circumventing the need for TCR/CD3 engagement. Activation was assessed through the expression of the activation marker CD69, revealing that both T cell clones could be activated (<80% WT T cell activation) but DN BAP1KO T cells exhibited compromised activation under ION-only stimulation conditions. (Fig. S5a-b). CD2 expression was assessed two days after stimulation and again at 11-12 days post-stimulation. As anticipated, WT Jurkat T cells exhibited a 1.8-fold increase in average CD2 expression upon stimulation compared to resting conditions, with similar increases observed for combinations of PMA with ION (1.2-1.8-fold) and high concentrations of ionomycin alone (1.5-fold); nearly all WT T cells were CD2+ across all conditions. In stimulated DN BAP1KO T cells, an upregulation of CD2 was detected, resulting in approximately 21% CD2+ DN T cells compared to 5% CD2+ in resting conditions (Fig. 6a). Despite this increase, CD2 levels in DN BAP1KO T cells remained significantly lower (about 8-fold less) than those found in resting WT T cells. Subsequent culture of T cell clones under resting conditions led to a reduction in the percentage of CD2+ DN BAP1KO T cells, except for those stimulated with high PMA concentrations, which maintained around 20% CD2+ DN T cells (Fig 6b, Fig S5c). This trend in CD2 expression contrasted sharply with the behavior of CD28 and CD226 upon stimulation followed by rest. While CD28 and CD226 were upregulated in WT T cells, they also showed significant upregulation in DN T cells under PMA-containing stimulations, reaching levels comparable to those in resting or activated WT T cells, respectively. Phenotypic analysis following subsequent culturing in resting conditions, showed that over 50% of DN T cells retained the upregulated CD28 or CD226 expression at levels similar to those observed in WT T cells (Fig. 6b). WT T cells strongly upregulated the co-inhibitory PD-1 receptor, upon high PMA-high ION stimulation (78% PD-1+ T cells) but DN BAP1KO T cells showed significantly lower percentage of PD-1+ T cells (42% PD-1+ T cells), despite similar levels of activation based on CD69 activation marker expression two days post-activation (Fig. S5b,c). Within PD-1+ T cells, the average PD-1 expression was lower in PD-1+ DN BAP1KO T cells (gMFI: 28851) compared to PD-1+ WT T cells (gMFI:70206). This suggests that like CD2 expression, BAP1 is required for efficient and high expression of PD-1 upon T cell activation. Finally, high ION with or without PMA stimulation resulted in increased TCR expression in both WT and DN BAP1KO T cells relative to resting condition (Fig 6a, Fig S5c). Despite this upregulation, DN BAP1KO T cells had a lower percentage of TCR+ T cells compared to WT T cells; in fact, average TCR expression was half-fold lower in TCR+ DN BAP1KO T cells compared to WT TCR+ T cells. Subsequent culturing in resting conditions resulted in a decrease of TCR+ T cells in both T cell clones; this contraction, though, was more pronounced in DN T cell population compared to WT T cells (Fig S5d-e). These findings reveal differential mechanisms governing the expression of costimulatory receptors upon T cell activation compared to resting conditions for distinct gene groups—namely CD2 and TCR versus CD28 and CD226—highlighting the context-dependent nature of gene regulation.

**Fig. 6.**
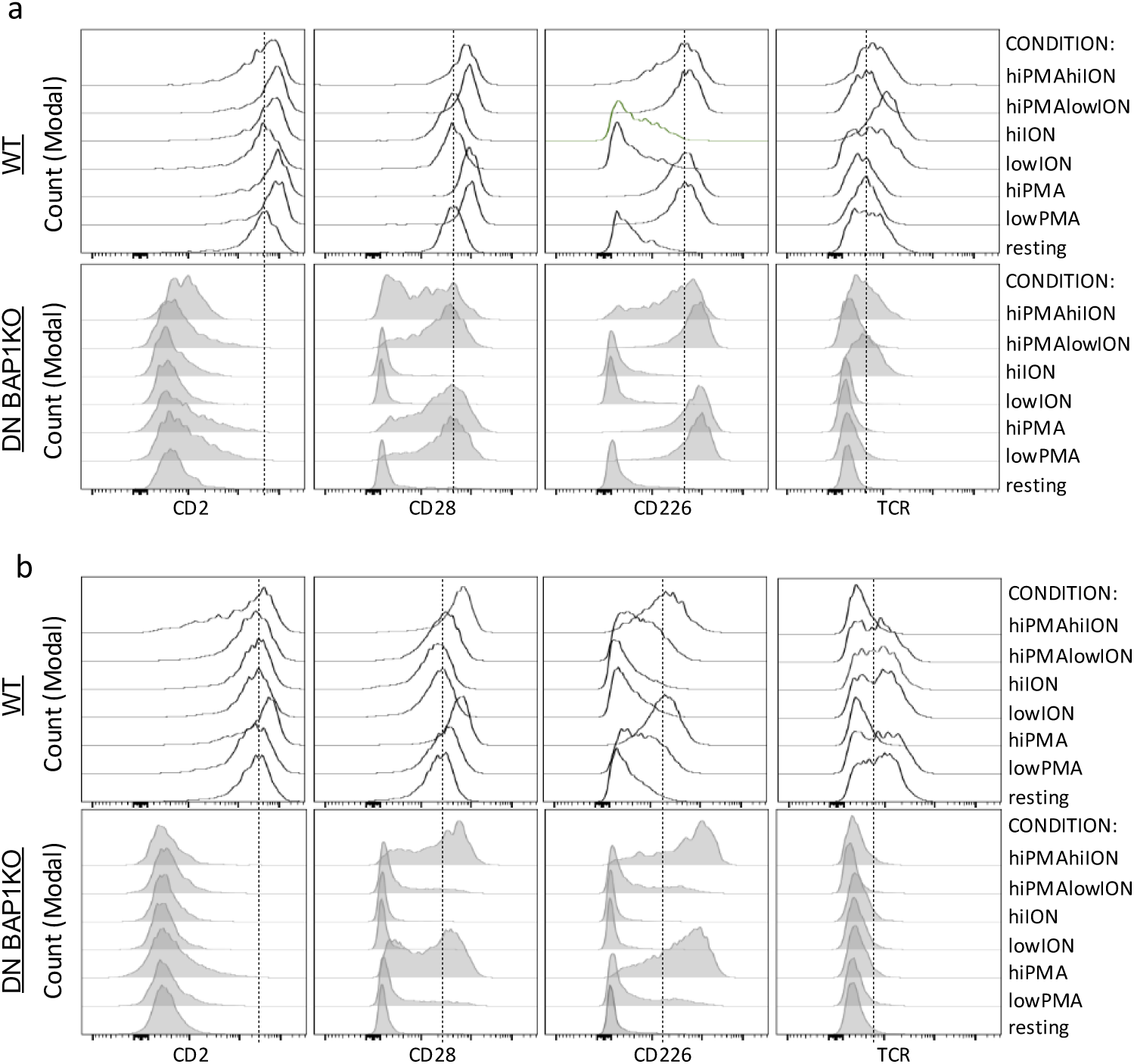
T cell activation reveals differential CD2 and TCR expression in BAP1 KO vs WT T cells but not for CD28 and CD226. **(a**) Flow cytometry histograms of CD2, CD28, CD226 and TCR expression in WT and DN BAP1KO T cell in resting conditions and two days post-chemical cell activation by-passing the need to engage the TCR-CD3 complex, with low (0.6μg/ml) and high (20μg/ml) concentrations of phorbol myristate acetate (PMA), with low (0.1μg/ml) and high ((1μg/ml)) concentrations of ionomycin (ION) or a combination of high concentration of PMA with low or high concentration of ION. (**b)** Same as in (a) but 11-12 days post-chemical activation that includes a resting period of cell cultured in no stimulation conditions

### Pharmacological inhibition of histone deacetylases partially rescues BAP1 loss-mediated CD2 repression

ATAC-seq analysis revealed that within and around the *CD2* locus there are condensed regions of chromatin relative to WT; this is similar for the *TOX* locus. This suggests that loss of BAP1 results in CD2 repression by through chromatin condensation that is one know way of transcriptional regulation^23,24^. One such epigenetic mechanism driving this chromatin condensation is the accumulation of tri-methylation of lysine 27 on histone H3 protein (H3K27me3) carried out by the Polycomb repressive complex 2^24,25^. We treated DN BAP1KO T cells with a number of PRC2 complex inhibitors (PRC2 catalytic subunit enhancer of zeste homolog 2, EZH2, inhibitors) to see if this rescues the loss of CD2 but there was no upregulation of CD2 (Fig. 7a, top). In contrast, EZH2 inhibitor treatment of WT cells resulted in a dose-dependent reduction of CD2 expression (Fig. S7b), agreeing with our GW CRISPR-Cas9 genetic screen that identified SUZ12 as a potential regulator of CD2 expression; SUZ12 is one of the members of the PRC2 complex together with EZH2. Repression of CD2 expression was confirmed in independent gene KOs of EZH2 (Fig. S7c). BAP1 KO has been shown to not only increase global H3K27me3 but also reduce oH3K27acetylation (K3K27ac) around gene promoters^26^. To investigate, whether counteracting loss of H3K27ac upon BAP1KO upregulates CD2 expression, we treated DN BAP1KO T cells with the histone deacetylase inhibitor, LMK235. This resulted in a partial rescue of CD2 loss upon BAP1 loss. There was a dose-dependent upregulation of CD2 expression with increasing levels of the HDAC inhibitor (Fig 7b); this upregulation though was not strong enough to reach the high levels seen in WT T cells. These findings suggest there is an interplay between BAP1 activity and HDAC activity in regulating CD2 expression. However, boosting H3K27ac is apparently not enough on its own to enable high levels of CD2 expression in the absence of BAP1.

**Fig. 7.**
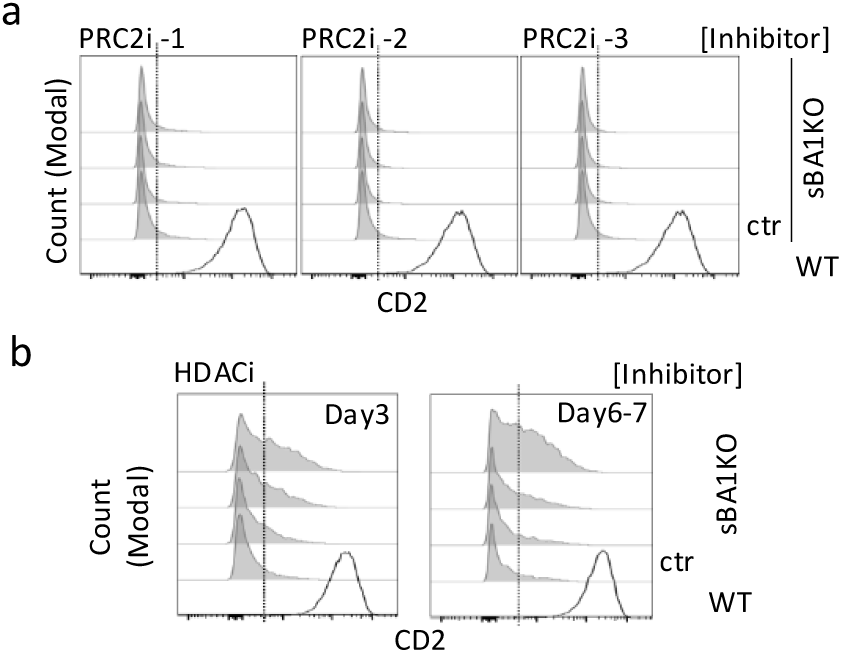
Pharmacological inhibition of histone deacetylases partially rescues BAP1 loss-mediated CD2 repression. **(a)** Flow cytometry histograms showing the surface CD2 expression in DN BAP1KO T cells in control condition and upon treatment with three different PRC2 inhibitors in a dose-dependent manner, seven days post-initiation of treatment (PRC2i-1, GSK126, PRC2i-2,A-395, PRC2i-3, Tazemetostat). **(b)** Flow cytometry histograms showing the surface CD2 expression in DN BAP1KO T cells in control condition and upon treatment with HDAC inhibitor, LMK235, in a dose-dependent manner, three and six/seven days post-initiation of treatment; a representative experiment shown from two independent ones. WT represent CD2 expression in WT T cells in control untreated conditions.

## DISCUSSION

Through, this work, we identified novel regulatory properties of the CD2-CD58 costimulatory pathway in human T cells; a pathway that has attracted significant interest in the fields of cancer immunology and immunotherapy^2,3,5,27–29^ over the past five years. Our findings reveal that CD2 costimulation strength regulates downstream T cell responses and identify positive regulators of CD2 expression, including the epigenetic regulators SUZ12 and BAP1. While focusing on BAP1, which has been understudied in T cell biology, we demonstrated that BAP1 acts as a broader regulator of key co-stimulatory and co-inhibitory receptors, beyond CD2, that are involved in T cell function and identity and it does so in a dose-dependent manner. Additionally, BAP1 co-regulated genes associated with T cell stemness, exhaustion, and metabolism.

We demonstrated that increasing CD2 costimulatory strength, not only enhances the percentage of T cells that get activated but it increases the average number of divisions (proliferation index, PI) and the fold expansion (replication index, RI) of these activated T cells. Furthermore, increasing CD2 costimulation strength correlated with increasing expression of the high affinity subunit of the IL-2 receptor, CD25, whose expression also shows a positive correlation with PI and RI. These correlations suggest that CD2 costimulation strength may regulate PI and RI by modulating CD25 expression levels, thereby influencing responses to the proliferation-inducing IL-2 cytokine. We also observed a similar titration effect of CD2 costimulation strength on IFN-γ production in an *in vitro* Th1 differentiation context. This aligns with a recent study showing that overexpression of CD58 in a melanoma cell line leads to higher IFN-γ production in an in vitro T cell:tumour cell co-culture system^30^.

Our results have significant implications for T cell anti-tumour immunity. We found that human TI-T cells in glioblastoma and meningioma, despite of their expression of memory markers, exhibit significantly reduced levels of CD2 resembling the levels observed in PB naïve T cells. This observation is reminiscent of the CD2^low^ phenotype we previously described in CD8+ TI-T cells in a percentage of CRC, EndoC and OC patients^5^. Moreover, in these GBM and MNG cohorts we also detected reduced CD2 expression in CD4+ TI-T cells in almost all patients; this was not exhibited by CD4+ TI-T cells in our previously studied cancer types and cohorts. This is important to consider since it has been shown that activation of both subsets is required for functional anti-tumour T cell immunity in brain tumours^14–16^ The reduction of CD2 expression in TI-T cell may serve as an immune evasion mechanism imposed by the tumour microenvironment, akin to the loss of CD58 observed in hematological malignancies^31,32^. This phenomenon could indicate a poor prognosis or predict inadequate responses to certain immunotherapies, similar to the lack of clinical benefit seen in melanoma patients with low CD58 expression when treated with immune checkpoint blockade^2^. In fact, in a recently published partially-humanised melanoma model study^2^, it has been shown that the absence of CD58 expression in tumour cells compromises T cell tumour infiltration, activation and cytotoxicity, similar to an earlier *in vitro* cytotoxicity screen^33^, further highlighting the potential impact reduced CD2 costimulation strength could have on human anti-tumour T cell immunity.

Through a genome-wide CRISPR-Cas9 KO screen, we report novel findings of genetic regulators of CD2 expression. We identified the epigenetic regulators SUZ12 and BAP1 as two positive regulators of CD2 expression. Focusing on BAP1 - given its limited characterization in T cells - and employing bulk RNA sequencing analysis, flow cytometry, gene ontology and GSE analyses, we determined that BAP1 acts as a broad regulator of gene expression in T cells. BAP1 KO affected the expression of a number of groups of genes including co-stimulatory/inhibitory receptors, beyond CD2, adhesion molecules, transcription factors involved in T cell development and differentiation, factors involved in the balance between T cell stemness and exhaustion and genes regulating cell metabolism. This agrees with the role of BAP1 described in other cell types especially in cancer cells as a regulator of gene expression by constraining ubiquitination of genome histones^34^ ^35^. We also show that BAP1 has a dose-dependent effect on the regulation of costimulatory receptors and stemness-exhaustion-associated factors. BAP1 overexpression studies such as this^36^ might have hinted on this dose-dependent effect before. Notably, BAP1 KO T cells exhibited a mixed phenotype characterized by enriched stemness and exhaustion markers. Further investigation is warranted to elucidate how the BAP1 KO-mediated expression changes in stemness- or exhaustion-associated factors influence downstream T cell anti-tumour immunity in vivo. Interestingly, a very recent study^37^ using a mouse chronic infection model and ASXL1 KO P14 T cells – ASXL1 is a key partner of BAP1 in the Polycomb repressive deubiquitinase complex, PR-DUB ^38,39^– showed that although ASXL1 KO T cells had higher frequency of Tex compared to WT T cells, bulk RNA sequencing of ASXL1 KO T cells showed lower expression of TOX, a Tex associated marker, in ASXL1 KO T cells and higher expression of Tpex associated genes such as *Tcf7* and *Myb*. Considering their findings and mechanistic insight describing that ASXL1 KO results in reduced activity of the PR-DUB complex, one might suggest that human BAP1 KO T cells might show a similar phenotype in an *in vivo* cancer setting but this is yet to be confirmed. However, a mouse study^40^ comparing BAP1 KO and ASXL1 KO tumour cells revealed significant similarities and differences in the genes regulated in each KO so one should take caution in extrapolations; the degree of similarity between the two KO phenotypes could be cell- and/or context-specific. Our transcriptomic findings showing that BAP1 regulates key factors involved in T cell development including the *TRAC* and pre-TCRA (*PTCRA*) and our GSEA findings that BAP1 KO has an enriched gene signature for genes downregulated in thymic T cell progenitors agree with the first study^41^ investigating mouse BAP1 KO T cells; it showed defective thymocyte development, impaired T cell proliferation in mature T cells in the periphery and impaired Th17 differentiation upon BAP1 KO.

Finally, we showed that BAP1 is essential for efficient high expression of CD2 upon T cell activation as CD2 was only partially upregulated and sustained upon T cell chemical activation in contrast to CD28 and CD226 that showed high and sustained expression. This suggests that T cell activation induces pathways that can rescue loss of CD28 and CD226 due to BAP1 KO, but these are not sufficient to restore CD2 expression. It is yet to be determined what are these exact pathways. CD2 expression though was rescued in BAP1 KO T cells by inhibiting histone deacetylases. A similar pharmacological rescue of a BAP1 KO phenotype was observed for the gastrulation process during embryo development of *Xenopus laevis*^26^. We suggest that this is the result of increasing histone acetylation that allows a more open chromatin structure^42,43^ counteracting the closer chromatin at the CD2 locus induced by BAP1 KO as shown by our ATAC-Seq analysis.

In summary, our work has contributed to three major advances in T cell biology. First, we elucidate a previously unappreciated role of CD2 costimulation strength in downstream T cell responses further boosting our suggestion that the CD2^low^ phenotype, now also described in glioblastoma and meningioma T cells, is another pathway impacting anti-tumour T cell immunity. Second, we identify positive regulators of CD2 expression including two epigenetic regulators, SUZ12 and BAP1. Finally, we demonstrate that under steady state conditions CD2 is co-regulated by BAP1 with other costimulatory receptors and stemness- and exhaustion-associated factors further highlighting the link of the CD2 pathway with other key pathways determining T cell responses in diseases such as cancer.

## MATERIALS AND METHODS

### Reagents

RPMI-1640 (31870074), Fetal Bovine Serum (FBS), Phosphate-buffered saline (PBS) and Dynabeads Human T-Activator CD3/ CD28 for T cell expansion and Activation kit (1132D) were purchased from Thermo Fisher Scientific. The following anti-human antibodies were purchased from Biolegend: CD62L (DREG-56), CCR7 (G043H7), CD3 (SK7), CD4 (SK3), CD8 (RPA-T8), CD28 (CD28.2), PD-1 (EH12.2H7), mouse IgG1 isotype control (MOPC-21), Human TruStain FcX, CD45 (HI30), CD45RA (HI100), CD2 (RPA2.10), CD57 (QA17A04), CD96 (NK92.39), CD11 (HI111), TCR-αβ (IP26), CD226 (11A8), CD127 (A019D5), CD49a (TS2/7), CD58 (TS2/9), CD3 (OKT3), CD69 (FN50), CD25 (M-A251). The following antibodies were purchased from Cell Signalling: BAP1 (D7W7O), CD3ε (D7A6E), TRBC1/TCRβ constant region 1 (E6Z3S), GAPDH (D4C6R), β-actin (8H10D10). The following antibody was purchased from Santa Cruz Biotechnology: TCR α Antibody (H-1). The following antibody was purchased from Proteintech: CD2 Monoclonal antibody (3A10B2). The following Taqman primer-probes were purchased from Thermo Fisher Scientific: Bap1 (Hs00184962_m1), Cd2 (Hs01040180_m1), Abl 1 (Hs01104728_m1), Tox (Hs01049519_m1), Tcf7 (Hs01556515_m1), Bach2 (Hs00935338), Id2 (Hs04187239). Antibodies were used at dilutions recommended by the manufacturer for western blotting and at concentrations in the range of 0.5-2 μg/ml for flow cytometry depending on the titration experiments and number of cells stained. The BlueStar Prestained Protein Marker (MWP03) was purchased from Nippon Genetics.

### Isolation of PBMCs and T cells from human samples

Human PBMCs and T cells were isolated from leukapheresis products of healthy controls (Cyprus National Bioethics Committee License 2020/44) or fresh peripheral blood of cancer patients (Cyprus National Bioethics Committee License 2020/45) using Ficoll-Paque density gradient as per suppliers instructions followed by an erythrocyte lysis step using ACK lysis buffer (ThermoFisher A1049201) as per manufacture’s protocol; T cell subsets were isolated using negative selection EasySep enrichment kits from STEM CELL Technologies. T cells were maintained when necessary in resting conditions, in the absence of any IL-2, at 37°C, 5% CO2, in RPMI medium (Life Technologies) supplemented with 10% FBS, 5% Penicillin-Streptomycin, (Penstrep, Gibco), 1x MEM Non-Essential Amino Acids Solution (Thermo Fisher Scientific), 10mM HEPES (Life Technologies), 1mM Sodium Pyruvate (Thermo Fisher Scientific). T cells were maintained at maximum 2×10^6^ cells/ml for a maximum of 5 days during resting conditions.

### Single cell suspensions of human brain tumour samples

Preparation of single cell suspensions from fresh brain tumor tissue (BT) was carried out as described previously in^44^. In brief, the sample was cut into small fragments (approximately 1– 2 mm^3^) and resuspended in Hanks’ Balanced Salt solution (HBSS^(+Ca+Mg)^; Sigma-Aldrich, cat. No. 55037C) at 100 mg tissue per ml. Then neutral protease (NP, AMSBio. S3030112) at 0.11 DMC U/ml and a Bovine pancreatic deoxyribonuclease I was used at 400U/ml (DNase I; Sigma-Aldrich, 11284932001) were added to the suspension and incubated for an average time of 2 hours at 37°C. The preparation was then filtered through a 70 μm strainer to remove clumps. The single cell suspension was then washed once and resuspended in phosphate-buffered saline (PBS, Fisher Scientific, 10010056) supplemented with 10% fetal bovine serum (FBS, ThermoFisher, A5256801). The standard trypan blue dye-exclusion method was used to evaluate cellular viability.

### Cancer patient cohort

Sixteen fresh tumour samples from two sides of the tumour and matched peripheral blood in EDTA-blood collection tubes from consented GBM and MNG patients were obtained from patients who underwent surgical removal of tumours at the Aretaieio Private Hospital (Nicosia, Cyprus). They were processed to generate single cell suspensions. Information about the age and gender of the patients is shown in Supplementary Table 3.

### Human primary T cell activation and proliferation assays

Proliferation of human T cells was monitored by following dilution of CFSE(Sigma-Aldrich)-labelled T cells (Sigma-Aldrich). Briefly, T cells single cell suspensions at 3×10^6^ cells/ml in PBS were stained with 1 μM CFSE in PBS solution by a 10 min incubation in a culture incubate at 37°C, 5% CO₂. Staining was quenched by adding FBS at a final 10% FBS concentration before pelleting the cells to removed supernatant. Cells were washed and resuspended in culture medium for downstream assays.Typically, T cells were prepared for T cell activation in U-bottom plates at 1×10^6^/ml and activated with anti-CD3/CD28 antibody coated Dynabeads for 2 days before removing the activation stimulations and supplementing the culture with 30U/ml IL-2. The culture was split every other day starting from the day of stimuli removal. At the endpoint, T cells were stained with fixable viability ef780 dye (ThermoFisher) followed by a mixture of fluorophore-conjugated antibodies when needed. Stained sampels were acquired on a 3-laser Aurora spectral flow cytometer (Cytek)

### Profiling human samples using flow cytometry

Human primary T cells, PBMCs, Jurkat T cells and cancer patient tumour single cell preparations were stained at around 0.05×10^6^, 0.01×10^6^, 0.1×10^6^ and 1×10^6^ per staining condition respectively in U-bottom plates. For PBMCs and cancer tumour samples an Fc-receptor blocking step was carried out by resuspending cells in 50 μL PBS with 0.5% BSA and 5 μL Human TruStain FcX (BioLegend) per sample, followed by incubation at room temperature for 10 minutes. Cells were stained with an antibody mix prepared in ice-cold PBS/0.5% BSA and incubated on ice in the dark for 20 minutes. Antibodies were used in the range of 0.5–2.5 µg/mL, based on the determined staining index of each antibody. Following the antibody staining, the cells were washed twice in PBS/0.5%BSA and acquired on a 3-laser Aurora flow cytometer. Data analysis was performed using FlowJo software (v10.8.1).

### In vitro differentiation of human naïve CD4+ T cells to Th1

Human naïve CD4+ T cells were isolated as described above. They were activated for 2 days using anti-CD3-CD28 antibody coated Dynabeads. The culture medium was supplemented with a cytokine-containing cocktail supplemented also with cytokine blocking antibodies from STEMCELL Technologies (ImmunoCult Human Th1 Differentiation Supplement, 10973) recommended for differentiation to Th1; it was used at 1 in 500 dilution. Following removal of activation beads, cells were split 1 in 2 every other day. Th1 status was profiled 7 days post-initiation of experiment using intracellular cytokine staining for IL-2 and IFN-γ. Intracellular cytokine staining was performed using the Foxp3 staining kit from eBioscience as per company’s instruction for a U-bottom two-step protocol.

### Genome wide (GW) CRISPR-Cas9 Knockout screen in Jurkat T cell line

For generation of constitutively Cas9-expressing cell lines, vector pKLV2-EF1a-Cas9Bsd-W was packaged into lentivirus by co-transfection of HEK293T cells alongside lentiviral envelope and packaging plasmids pMD2.G and psPAX2 using Lipofectamine LTX with PLUS reagent (Invitrogen) as described in^45^. The Jurkat cell line was transduced with lentivirus carrying Cas9 in the presence of polybrene (8 μg/ml). The following day, lentiviral containing medium was replaced with complete medium. Blasticidin selection was started on day 4 post transduction at 4 μg/ml. Cas9 activity was assessed using a lentiviral reporter system consisting of the constructs pKLV2-U6gRNA(gGFP)-PGKBFP2AGFP-W and pKLV2-U6gRNA(gEmpty)-PGKBFP2AGFP-W. Cells were transduced with a predetermined viral amount of the genome wide gRNA-BFP library that allows for an average of 30% transduction, measured by BFP expression by cytometry and ensures incorporation of a sgRNA per cell with about 9% error rate. Transduction was performed at 1×10^6^ cells per well in 6-well plates for a total of 100×10^6^ to allow for a 350x coverage of the library. Puromycin selection was used, starting at day 4 of the culture, to select for cells that had successful lentiviral plasmid integration. Cell transduction efficiency was determined before and after puromycin selection by quantifying the percentage of BFP expressing cells using flow cytometry. FACSorting was performed at 5 different timepoints during cell culture of these cells, sorting for 5% of CD2low/neg cells from the live T cell population. FACSorted T cells were stored in RNAlater in −20°C till DNA extraction, DNA library preparation and next-generation sequencing. The bioinformatic analysis was performed as described previously in^44,46^ using the MaGeCK package.

### Designing single guide RNAs(sgRNA) for sgRNA knock outs in T cells

sgRNAs were designed using the CHOPCHOP and Synthego web tool or using the gRNA list from the Human genome-wide lentiviral CRISPR gRNA library version 1(Addgene 67989) (Supplementary Table 4). Priority was given to plasmids found enriched in the MaGeCK analysis results of our screen. For online tools, selection criteria also included GC content (>45%), efficiency score (>0.5), and minimal self-complementarity/mismatch scores.

### Western blotting

Whole cell lysates were prepared by incubation of the cell pellet for 30 mins in 1X RIPA buffer (Sigma-Aldrich, R0278) supplemented with 1 mM PMSF and 1X protease inhibitor cocktail (Sigma-Aldrich, 11836153001) on ice. Supernatant was harvested and stored in -20°C till required for downstream analysis. Total protein concentration was determined using the Bradford assay. 50–100 µg of total protein was loaded per well in a 8 or 10% SDS-PAGE gel depending on the molecular weight of the proteins of interest. Whole cell lysates were prepared using 1X Laemmli loading buffer (Bio-Rad, 1610747) and denatured for 5 mins at 95°C. Proteins were transferred to Porablot nitrocellulose membranes (Macherey-Nagel, 741280) using a wet transfer system. Membranes were blocked in 5% nonfat milk or BSA prepared in in TBS-T (Tris-buffered saline supplemented with 0.05% Tween-20) for 1 hour at room temperature and incubated overnight at 4 °C with primary antibodies prepared in TBS-T supplanted with 5%BSA or milk as per manufacturer’s recommendations. After washing with TBS-T, membranes were incubated with a IRD680RD or IRD800CW-conjugated secondary antibody for 1 hour at room temperature. Following extensive washing, protein bands were imaged on the Vilber imaging system. Protein band total intensities were quantified using FIJI software, normalized to housekeeping proteins (β-actin or GAPDH) and presented as relative protein expression.

### RNA extraction, real-time quantitative PCR, bulk RNA sequencing and differential expression gene analyses

Total RNA was extracted using the MagCore® triXact RNA Kit (Cat.no. 631) as per manufacturer’s recommendations and assessed for quality and quantity using a NanoDrop™ 2000/2000c Spectrophotometer. The RNA aliquots were stored at −80°C.

For real-time quantitative PCR (RT-qPCR), total RNA from each sample was used to synthesize cDNA using PrimeScript RT Reagent Kit(Takara, cat. No RR037A) according to the manufacturer’s protocol. The reagents, RNA, oligo (dT), and dNTPs were mixed first, then incubated at 37 °C for 15 mins, 85 °C for 5 mins and then chilled on ice. The samples were stored at -20°C till required for downstream assays. RT-qPCR was performed using the TaqMan™ Universal PCR Master Mix (Thermo Fisher Scientific, 4304437) for precise quantification of gene expression. TaqMan primer-FAM probe pairs were purchased from Thermo Fisher Scientific and are included in the reagents section above. The internal control gene used was *ABL1* (Hs01104728_m1), and target genes used can be found in the reagents paragraph. All of the reactions were run in triplicates on a CFX 96 (BioRad) thermal cycler. The Cq values were estimated by analysing the data as a single pool, using the automatic threshold and baseline cycle option of the CFX Manager Software v3.1. Amplifications were conducted in triplicates on a real-time PCR system under the following thermal cycling conditions: an initial step at 50 °C for 2 minutes, 95 °C for 10 minutes, followed by 50 cycles of 95 °C for 15 seconds and 60 °C for 1 minute. Fluorescence was monitored throughout amplification. Relative gene expression levels were calculated using the 2^^−ΔΔCT^ method, normalized to *ABL1*.

cDNA libraries for bulk RNAsequening analysis were prepared from the extracted total RNA using the QuantSeq 3’mRNA-Seq V2 Library Prep Kit-FWD (Lexogen; 192.96) according to the manufacturer’s guidelines. Three biological replicates of each sample were analyzed. The pooled libraries were sequenced on an Illumina the NextSeq2000 system. Different expression gene analysis was performed by the Lexogen pipeline using DESeq2 algorithm as described on the company’s website.

### Bioinformatic analysis including gene ontology, KEGG and Gene Set Enrichment Analyses

Bioinformatics analysis of differential expression findings was performed by utilizing the R package clusterProfiler for functional enrichment analysis. Gene Ontology (GO) and Kyoto Encyclopedia of Genes and Genomes (KEGG) enrichment analyses were conducted to identify over-represented biological themes. Additionally, Gene Set Enrichment Analysis (GSEA) was performed using gene sets from the Molecular Signatures Database (MSigDB) collections C5 (ontology gene sets) and C7 (immunologic gene sets). All analyses were conducted using default parameters within the respective functions. For the generation of a Tpex signature, we utilised bulk RNAseq data from ^8^ and comparison of CD101-TIM-3-T cells and CD101+TIM-3+ using the DESeq2 software package; differentially expessed genes with a padj<0.05 and log2fold change>0 were used as a Tpex gene signature against which a GSEA was performed with the DEG of BAP1 KO Jurkat T cells.

### Assay for Transposase-Accessible Chromatin using sequencing (ATAC-Seq) analysis

ATAC-seq analysis was performed using commercially available kit (Activemotif, 53150,) for cell processing and library preparation. In brief, 50,000 cells were processed using the above kit with one deviation of the incubation time required for the Tn5-mediated tagmentation of the DNA that was extended to 1 hr instead of 30 mins as recommended in the manual of the kit. Sample quality was assessed on an Agilent DNA tapestation using an Agilent D1000 screentape. Sequencing was performed on a NextSeq 2000 as per company’s instructions and analysis was performed by the the ATAC-seq analysis service of ActiveMotif company.

### Phorbol 12-myristate 12-acetate (PMA) and ionomycin treatment of T cells

Wild Type (WT) or FACSorted CD3-CD2-(double negative, DN) Jurkat T cell subsets were cultured in RPMI 1640 medium supplemented with 10% FBS, 2 mM L-glutamine, 100 IU/mL penicillin, and 100 μg/mL streptomycin in a 37°C incubator with 5% CO₂ with or without treatment with low levels of PMA (0.6ng/ml), high levels of PMA (20ng/ml) low levels of Ionomycin (ION, 0.1μg/ml), high levels of ION, 1μg/ml or high levels of PMA combined to low or high concentration of ION. Following two days of T cell treatment, culture media was replaced and cells split every other day in fresh supplemented media in resting, no stimulation conditions for an additional 9-10 days. T cell viability and expression of surface receptors was evaluated 2 days post-stimulation and after 9-10 in resting culture conditions using flow cytometry analysis.

### PRC2 and HDAC inhibitors treatment of Jurkat T cell subsets

Wild Type (WT) or FACSorted CD3-CD2-(double negative, DN) Jurkat T cell subsets were cultured in supplemented RPMI 1640 medium with or without PRC2 complex inhibitors (MedChemExpress, A-395, HY-101512; GSK126, HY-13470; Tazemetostat, Cat. No.: HY-13803). Inhibitor treatment, at titrated concentrations, was started on day 0 and supplemented on day 3 post-initiation of experiment. T cell viability and CD2 expression were evaluated 3 and 7 days post-initiation of experiment.

### Data analysis and statistics

Data were analysed by GraphPad Prism v10 wherever needed unless mentioned otherwise such as in the bioinformatics sections of bulk RNAseq analysis.

## ACKNOWLEDGEMENTS

We wish to thank Dimitris Aspris for his assistance in transfering the knowledge on CRISPR-Cas9 genome wide screens in cell lines to CSHM, Skevi Charalampous and all the nursing staff of Aretaieio Hospital, Nicosia for helping out with the patient consents and sample acquisition, respectively.

## Author contributions

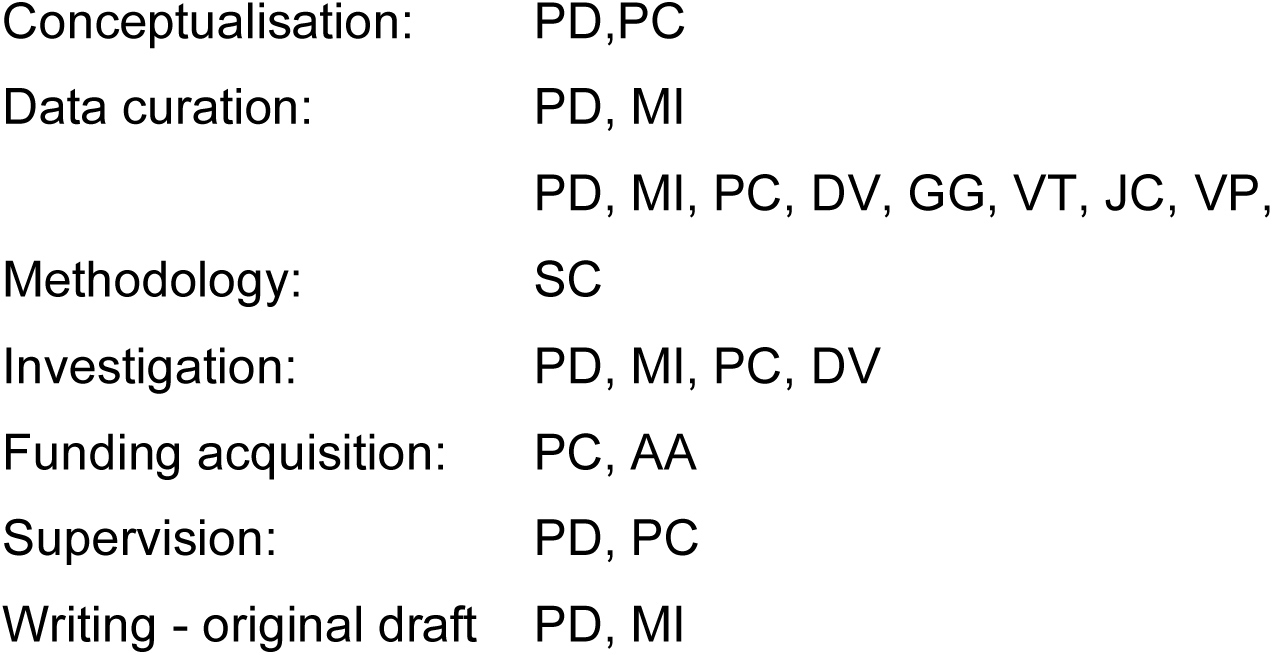

## DATA AND MATERIALS AVAILABILITY

The data that support the findings of this study are available from the corresponding author upon reasonable request.

