## Supplementary Figures for "CD2 expression is co-regulated with stemness- and exhaustion-associated factors in human T cells"

Supplementary Information

**Supplementary Fig 1. T cell proliferation and IFN-γ production correlate with CD2 costimulation strength**

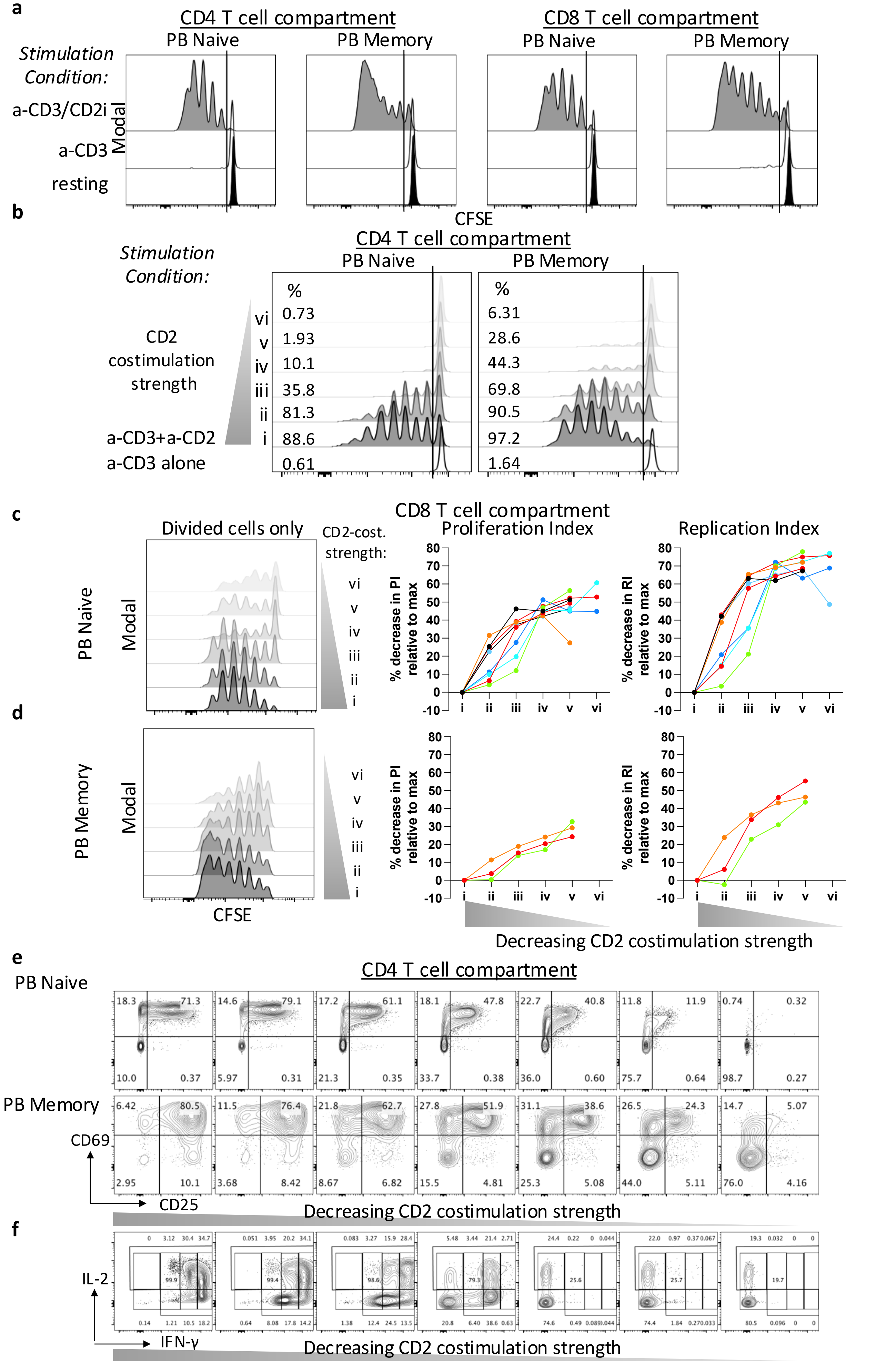

**Supplementary Fig.1. T cell proliferation and IFN-γ production correlate with CD2 costimulation strength. a-b)** Representative flow cytometry histograms showing gating strategy for non-divided and divided cells across conditions (resting i.e. no activation, anti-CD3 antibody (Ab) alone coated activation beads, a-CD3, anti-CD3 and anti-CD2 Ab coated beads, a-CD3/CD2); i)-vi) represent anti-CD3 Ab-coated beads with decreasing levels of anti-CD2 Ab. Representative histograms (left) from one healthy volunteer, illustrating cell divisions of PB CFSE-labelled naïve **(c)** and memory **(d)** CD8+ T cells, gated on divided cells and corresponding plots (right) of the percentage decrease in proliferation index and replication index at titrated levels of CD2 costimulation strength (i-vi), 5-days post *in-vitro* activation from 8 and 3 donors respectively (each a different colour). **(e)** Representative dot plots of CD69-CD25 expression profile related to Fig. 1. c-d used to track % of activated cells across titrated levels of CD2 costimulation strength. **(f)** Representative dot plots of IL-2-IFN-γ expression profile related to Fig. 1. e-f used to track % of cytokine producing cells across titrated levels of CD2 costimulation strength 7 days post-activation and Th1 differentiation.

#### Supplementary Fig. 2. Naïve and memory profiles in peripheral blood and tumours of brain cancer patients

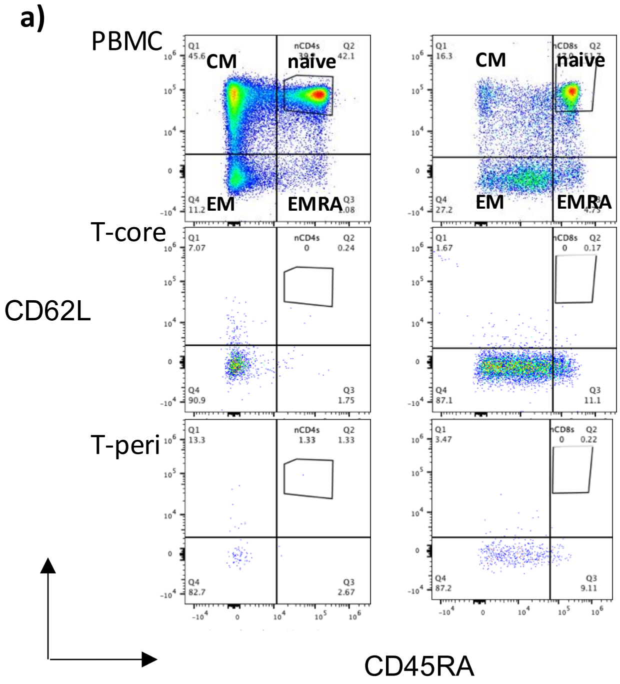

**Supplementary Fig. 2. Naïve and memory profiles in peripheral blood and tumours of brain cancer patients.** (a) Flow cytometry dot plots showing representative CD62L and CD45RA expression profiles and gates in peripheral blood mononuclear cells (PBMCs), TI-T cells from core and peripheral sides of tumour used to categorise in broad naïve, central memory (CM), effector memory (EM) and effector-memory re-expressing CD45RA (EMRA) T cell subsets.

**Supplementary Fig. 3. CD2 expression is coregulated with expression of markers associated with T cell development, stemness, exhaustion and metabolism**

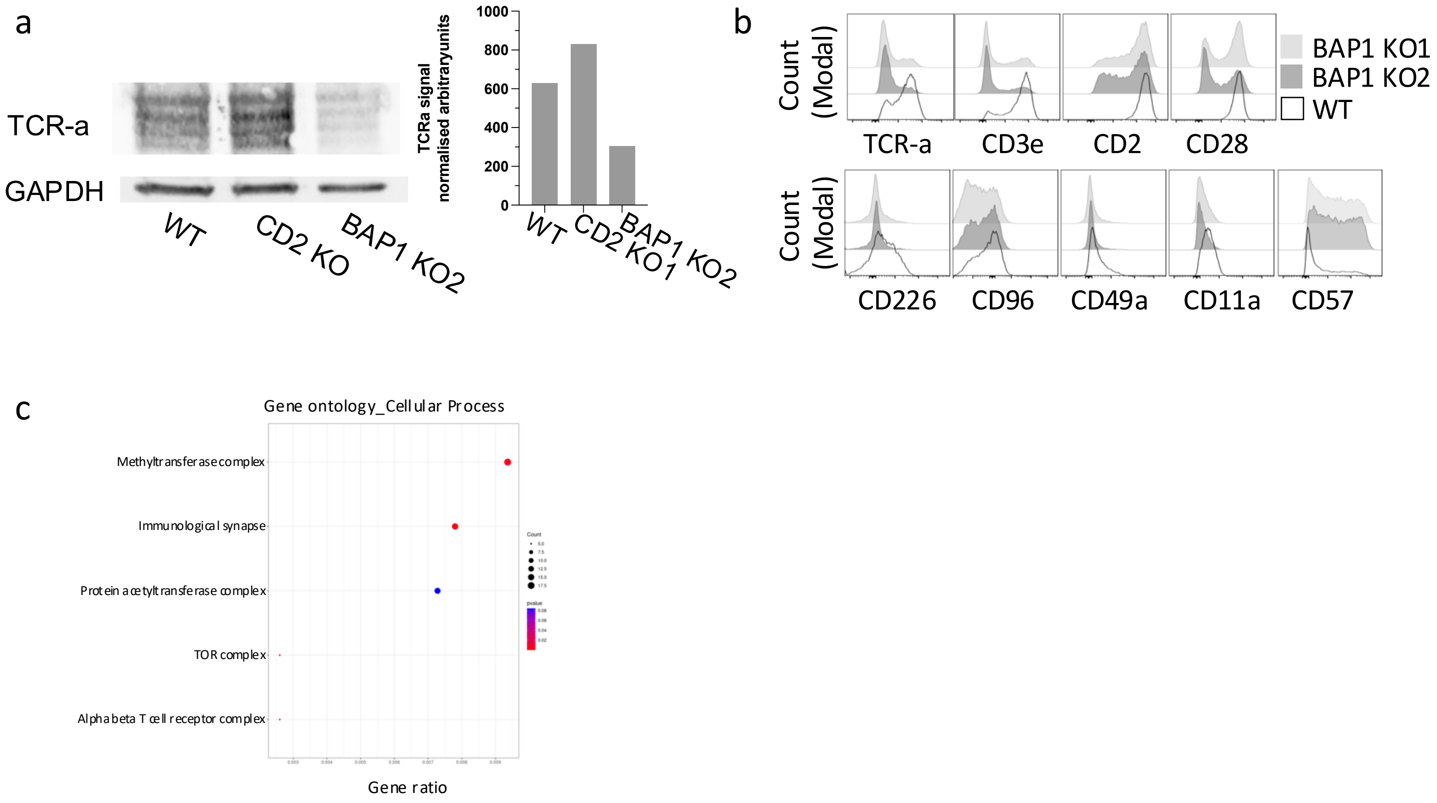

**Supplementary Fig. 3. CD2 expression is coregulated with expression of markers associated with T cell development, stemness, exhaustion and metabolism. (a)** Flow cytometry histograms showing the protein surface expression receptors in T cells upon BAP1 KO using two different sgRNAs compared to WT control; related to Fig.4b representative of two independent staining experiments and timepoints. **(b)** Western blot and corresponding bar chart showing the total amount of TCR-a protein normalized to b-actin expression in experiments as in Fig. 3d. **(c)** Bubble plot showing a selection of enriched pathways determined by performing gene ontology analysis (cellular process pathways) of DEGs in BAP1 KO T cells with a padjust<0.05. The gene count per pathway is represented by the size of the dot and the p value by the colour of the dot.

**Supplementary Fig. 4. BAP1 has dose-dependent effect on costimulatory receptor expression and stemness-exhaustion transcription factors**

a

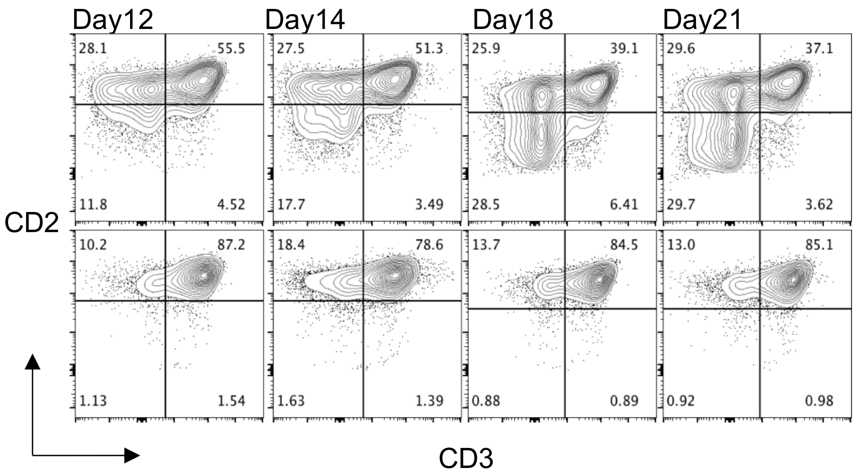


b

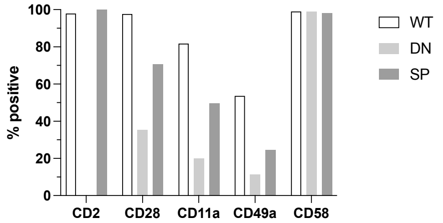

**Supplementary Fig.4. BAP1 has dose-dependent effect on costimulatory receptor expression and stemness-exhaustion transcription factors. (a)** Flow cytometry contour plots showing CD2-CD3e expression profile during sgRNA BAP1 KO1 in Jurkat T cells at different timepoints of the culture and the gating to define different populations, double positive, DP: CD3+CD2+, single positive, SP: CD3-CD2+/high, double negative DN: CD3-CD2-/low.) **(b)** Bar charts showing the percentage positive T cells for CD2, CD28, CD11a, CD49a, CD58 in DN, SP BAP1KO1 T cell subsets and WT T cells.

### Supplementary Fig. 5. T cell activation reveals differential CD2 and TCR expression in WT versus BAP1 KO T Cells but not for CD28 and CD226.

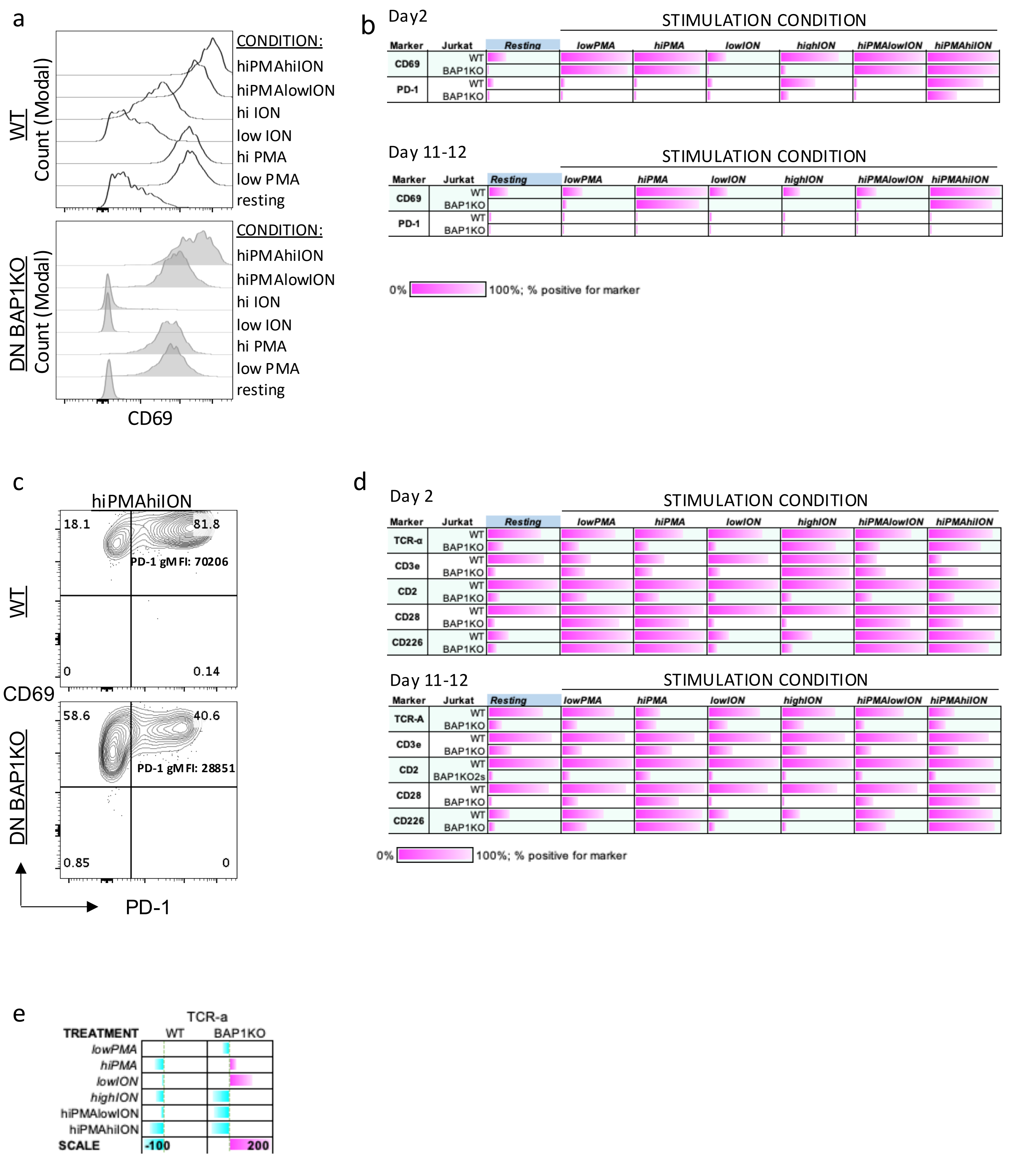

**Supplementary Fig. 5. T cell activation reveals differential CD2 and TCR expression in BAP1 KO vs WT T cells but not for CD28 and CD226. (a)** Flow cytometry histograms of the activation marker CD69 in WT and DN BAP1KO T cell in resting conditions and two days post-chemical cell activation with two different concentrations of PMA, ION or combination of the two agents; related to Fig. 6a. **(b)** Bar charts showing the % of positive cells for CD69 and PD-1 expression in DN BAP1 KO and WT T cells in resting condition and upon different stimulation conditions two days post-stimulation and 11-12 days post-stimulation; related to Fig 6a-b. (c) Flow cytometry dot plots showing the CD69-PD-1 expression profile in hiPMAhiION conditions in WT and DN BAP1 KO T cells two days post-stimulation.

### Supplementary Fig. 6. Pharmacological and genetic inhibition of EZH2 results in reduced CD2 expression in WT T cells

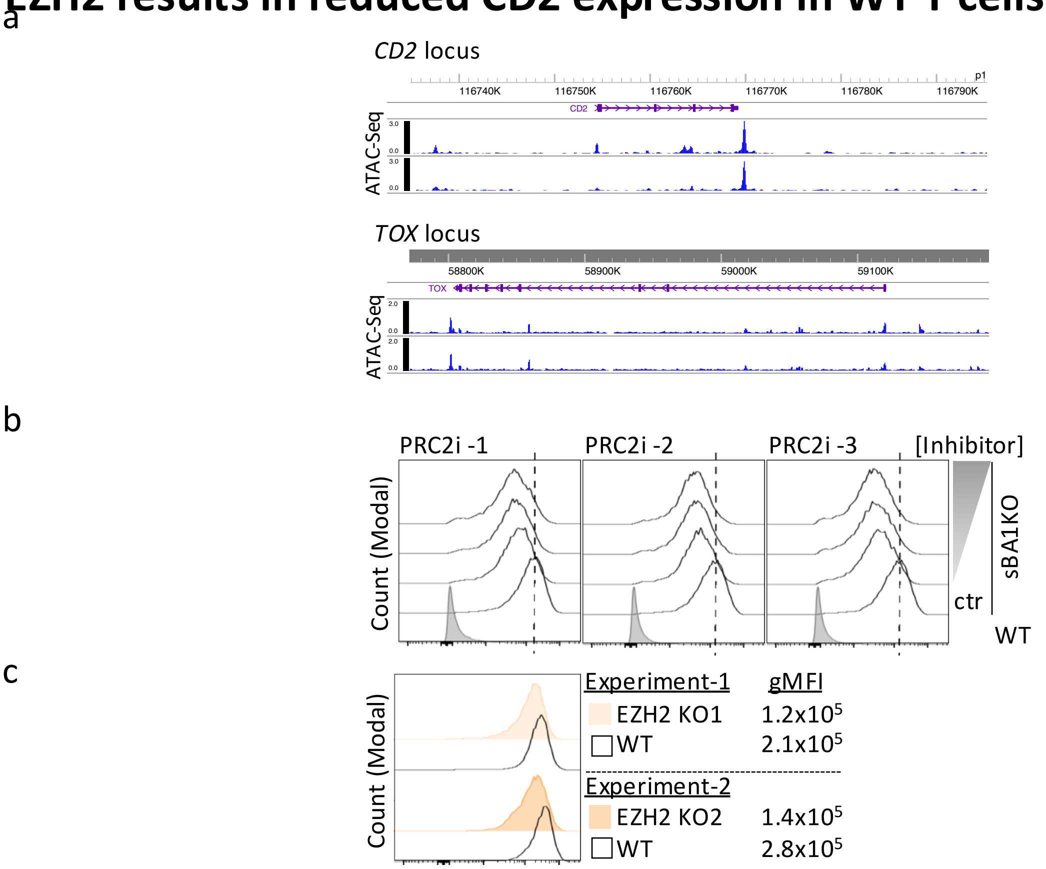

**Supplementary Fig.6. Pharmacological and genetic inhibition of EZH2 results in reduced CD2 expression in WT T cells. (a)** ATAC-seq analysis histograms of the CD2 and TOX loci in WT and DN BAP1KO T cells showing differential accessible regions highlighted in yellow. **(b)** Flow cytometry histograms showing the surface CD2 expression in WT T cells in control conditions and upon treatment with three different PRC2 inhibitors (A-395, HY-101512; GSK126, HY-13470 ; Tazemetostat , HY-13803 ) in a dose-dependent manner, seven days post-initiation of treatment. **(c)** Flow cytometry histograms showing the surface CD2 expression 3 weeks post EZH2 KO using two different sgRNAs targeting EZH2 (KO1 and KO2) performed in two independent experiments
